# Activity of acetic acid and sodium bicarbonate to inhibit *Asergillus ochraceus* on culture media and detoxification of ochratoxin A in poultry diet

**DOI:** 10.1101/2020.10.25.268177

**Authors:** Halima Z. Hussein, Mneer Saed Al-Baldawy, Rakib. A. Hamed

## Abstract

The study was conducted to evaluate the activity of acetic acid and sodium bicarbonate to inhibit *Aspergillus ochraceus* on culture media and reduce ochra toxin A in poultry diet. It was found that addition of acetic acid into PDA medium 0.2%, 0.5% and 4% could reduce A *ochraceus* growth by 90.22%, 98.77and 100%, respectively. Similar reduction in *A ochraceus* growth was obtained on PDA containing sodium bicarbonate that attained to 81%, 91.5%, and 100% at 4%, 2% and 5% concentrations, respectively. Results showed that addition of acetic acid at 0.5% and sodium bicarbonate at 1% could reduce ochre A mycotoxin by 65.7% and 43.6%, respectively in the first month and then reached to 100% in the second month for the two compounds.

## Introduction

Corn is one of the most important cereal crops worldwide as it is consumed by human and forms most poultry and animal diets (Abbas and Shier, 2009). In Iraq, cereals in general occupy most of the agricultural cultivated lands (FAOstats, 2019); and as a result, the spread of different pathogens affects the yield negatively (Al-Ani et al., 2011, 2012). It has been reported that corn seeds infected with many fungi producing mycotoxin including *A. ochraccus* that produce ochra A mycotoxin under field and storage conditions (Halima et al, 2015). The corn seeds contaminated with mycotoxins is a very serious problem for human, poultry and animals (Abbas and Shier, 2009, Eman, 2011, FAO,2011, Meulenberg, 2012, Hassan et al, 2018).

Several methods were adopted to prevent the contamination of corn seeds and diets with mycotoxins or mycotoxin degradation, among them is the use of chemical products. Sodium bicarbonate, sodium hydroxide, ammonia, hydrogen peroxide, formaldehyde, ammonium phosphate, nanoparticles, many acids and bases were widely used to prevent mycotoxins production and degradation in the diets (Paskevicius et al, 2006, Scott and Turcksecs, 2009, Frcitas - Silva and Venancia, 2010, Eman, 2011, Karlovsky, 2011, Husseni et al, 2017, Hussein et al 2018). The treatment of *ochre* A contaminated corn seeds with formic acid, propionic acid and formic acid of 0.25-1% for 3-24 hrs caused the degradation of the mycotoxin (Eman, 2011). This study was conducted to evaluate the efficacy of acetic acid and sodium bicarbonate to inhibit *Aspergilus ochraceus* growth and degrade *ochre A* mycotoxin on culture media and in poultry diet.

## Materials and methods

### Ochra A toxin extraction

An isolate of *A. ochraceus* producing ochre toxin A was obtained from Mycotoxin Laboratory, Plant Protection Department, College of Agriculture Engineering Sciences, University of Baghdad. This isolate was grown on liquid yeast extract medium and the isolate suspension was filtered through Whatman no. 4 filter paper. One hundred ml of chloroform was added to 50 ml of the filtrate in 250ml separating funnel. The mixture was agitated and let for 15 min to settle. The lower layer containing chloroform was passed through Whatman filter paper containing Na_2_SO_4_ to eliminate water and the filtrate was dried in water bath at 50 c. The dry extract was dissolved in 5 ml of chloroform and maintained under freezing.

### Ochra toxin A detection

Ten microliters of extract were spotted on thin layer chromatography (TLC) plate (20*20 cm) at 2 cm spacing, pre-activated for one hour at 110 °**C**. A spot of standard ochre toxin A was added at the plate left side for control. The plate was placed in glass basin containing a mixture of toluene: ethyl acetate: formic acid (60:30:10) at depth of 1cm. The plate was removed from the basin when the solvent reached to 2m from the upper side. The plate was let to dry at room temperature and tested under UV light at 366 nm using UV viewing cabinet (Asensio et al, 1982).

### Ochra toxin production

*A. ochraceus* isolate was grown on rice seeds. Hundred and fifty g of rice seeds were added to 100 ml dist. water in glass plate and autoclaved at 121°**C**, 1.5 kg /cm^2^ for 20 mins in two successive days. The seeds were inoculated with two 9 mm discs sliced from *A. ochraceus* isolate culture on PDA. The plate was well agitated for homogenization and maintained at 25 °**C** for 21 days. The contaminated seeds were oven dried at 50 °**C** in paper sacks and ground (Abbas at al., 1984). The concentration of ochra toxin A in corn seeds powder was estimated by HPLC at Ministry of Science and Technology as described by Caprita et al (2007).

### Activity of acetic acid and sodium bicarbonate to inhibit *Aspergillus ochraceus* growth

Acetic acid and sodium bicarbonate were added to PDA medium before solidification at 3 concentrations, 0.2, 0.5 and 1% and 1, 2 and 5 % respectively, (at 40-45 °**C**) and poured in petri plate of 9 cm diameter. The plates were inoculated with 7 mm discs from isolate culture (1 disc / plate) and maintained at 25°**C** for two days. The inhibition percentages were calculated using the formula:

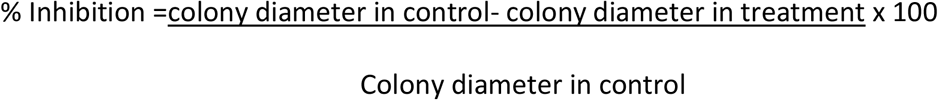

### Activity of acetic acid and sodium bicarbonate to reduce Ochra Toxin A in poultry diet

The experiment was conducted in Mycotoxin Lab. at Plant Protection Dept. for two months (1/4/2016 – 1/6/2018).

Fifty hundred g of poultry diet were placed in a desiccator and a quantity of consummated rice seed powder was added in to the diet to obtain 2mg / kg of ochra A.

Acetic acid at 0.5 % and sodium bicarbonate at 1% was added in to the contaminated diet and well agitated for homogenization. The treatments were randomized to three replicates and the concentrations of ochra A in the diet were detected on regular basis by HPLC.

## Results

### Activity of acetic acid and sodium bicarbonate to Inhibit *A. ochraceus* growth in PDA

It was found that addition of acetic acid in to PDA medium caused high gowth reduction of *A. ochraceus* ranged 90.22% −98.77% at 0.2% and 0.5%, respectively, while the concentration 1% inhibited fungus growth up to 100% (table 1 and fig1).

**Table (1):** Activity of different concentrations of acetic acid and sodium bicarbonate in Inhibition *Aspergillus ochraceus*

| Products | Concertation | Growth inhibition% |
| --- | --- | --- |
| Acetic acid | 0.2 | 90 – 22 |
|  | 0.5 | 90 – 77 |
|  | 1 | 100 |
| Sodium bicarbonate | 1 | 81 -50 |
|  | 2 | 100 |
|  | 5 | 100 |
| LSD P= 0.05 |  | 9.64 |

**Fig (1):**
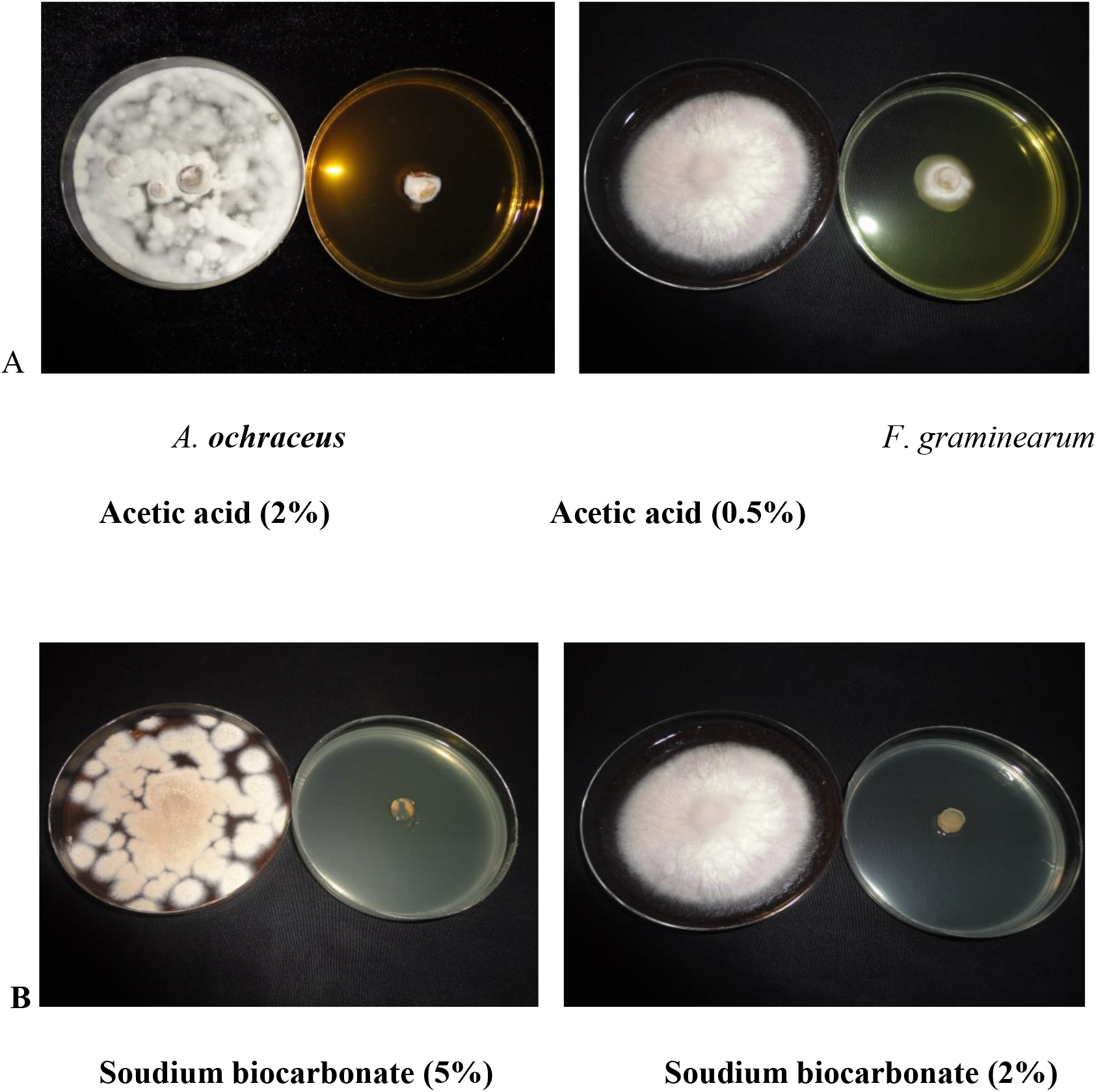
Effect of different concentration of Acetic acid and sodium bicarbonate on *A. ochracies* growth

Similar growth reduction in *A. ochraceus* growth was obsarved on PDA containing sodium bicarbonate which was 81% and 91.50 % at 1% and 2%, respectively compared to 100% at 5% concentration (table 1 and fig 1).

### Activity of acetic acid and sodium bicarbonate in ochra toxin A detoxification in the poultry diet

Results shown in table (2), revealed that the addition of acetic acid in to poultry diet at 0.5%could decrease ochra A mycotoxin concentration by 879.9 ng/g in the first month compared with 2566.1 ng /g in the control. at 65.7%reduction percent. Wherease, sodium bicarbonate at 1% reduced ochra A concentrations in the treated diet when scored 1446.4 ng /g in the first month compared with 2566.1 ng / g in control treatment with 43.6 %. The reduction of ochra A was 100% in the second month for two compounds as shown in table (2), fig (2).

**Table (2):**
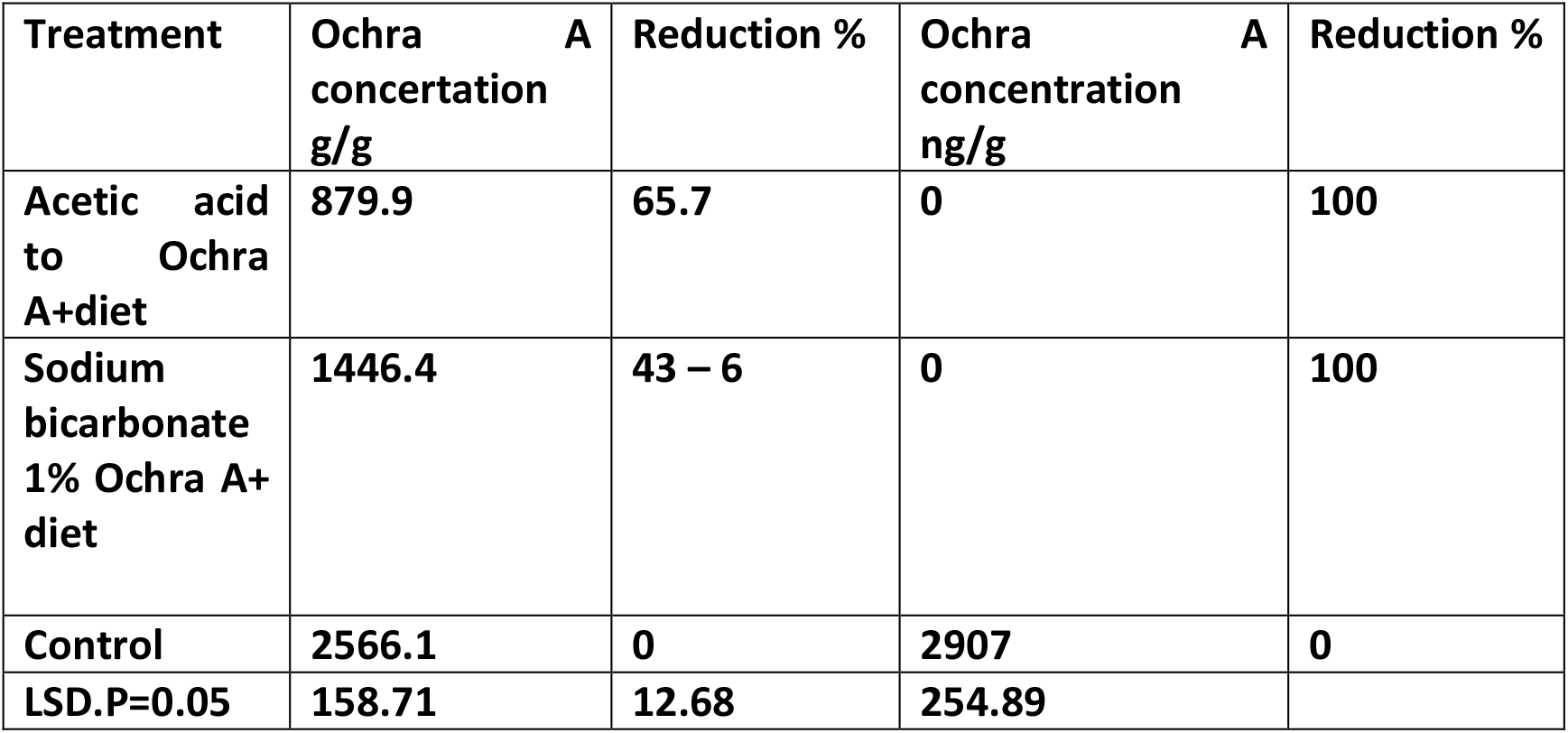
Efficacy of acetic acid and sodium bicarbonate in reduction ochratoxin A in poultry diets.

**Fig(2):**
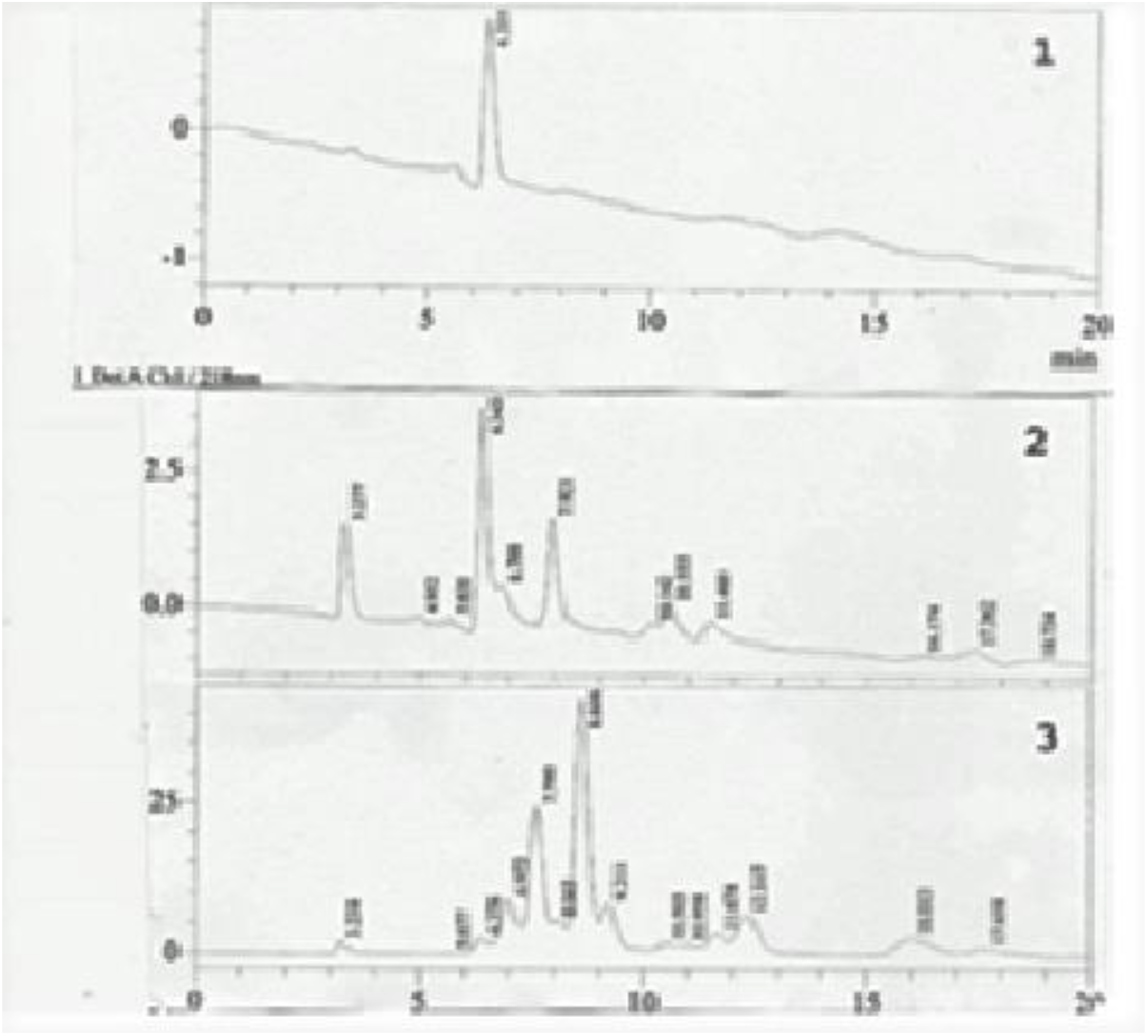
Activity of acetic acid and sodium bicarbonate in ochra A degradation in poultry diets

## Discussion

The results of this study have demonstrated the activity of acetic acid and sodium bicarbonate to inhibit *A*. *ochraceus* growth on culture media and their efficacy to degrade ochra A mycotoxin in poultry diet.

The activity of these two compounds in fungus growth inhibition may come from their ability to penetrate fungus cells and react with cell membranes causing deformation in the structure and modification in permeability. The interfering of these compounds with the membrane may result in fungus cell death.

It was reported that many acids including acetic and propionic acids showed high activity to inhibit the growth of fungi infecting corn seeds on culture media (Riess,1976, Moerch et al,1980, Mashely et al, 1983, Ali et al, 2011).

The activity of acetic acid and sodium bicarbonate in degrading ochra A may be attributed to the ability of these compounds to react with the active groups in the mycotoxins resulting in it’s the degradation or reducing their toxicity. Several studies have reported acid activity in the inhibition of A. ochraceus in stored seeds. Reiss (1976) found that addition of citric acid at 0.5 % and lactic acid at 0.75% inhibited *A. parasitica* growth in bread wheat. Juber and AL-salahi (2005) found that acetic acid at 2% and propionic acid at 10.5% were highly efficient to inhibit *A*. *flavus* and *A*. *parasitica* growth in stored wheat grains.

It was reported that the action of sodium bicarbonate in ochra A is on the lactone ring leading to break this ring and producing two type of products, (OP-OA) and (OA-OH) that have lower toxicity than Ochra A (Li et al, 2001). These results are in accordance with other studies that indicated the activity of bases in degradation of mycotoxins (AL-nazal, 2004). The addition of urea at 7% in to the diet contaminated with AFB1 (2mg/kg) caused mycotoxin degradation the (Majeed,1997)

